# Peripheral serotonergic neurons regulate gut motility and anxiety-like behavior

**DOI:** 10.1101/2023.05.12.540588

**Authors:** Hailong Zhang, Deborah R. Leitner, Yuko Hasegawa, Matthew K. Waldor

## Abstract

Serotonergic circuits in the central nervous system play important roles in regulating mood and behavior, yet the functions of peripheral serotonergic neurons are less understood. Here, we engineered mice lacking the serotonin-producing enzyme *Tph2* in peripheral neurons but with intact *Tph2* in central neurons. In contrast to mice lacking *Tph2* in all neurons, mice lacking *Tph2* in peripheral serotonergic neurons did not exhibit increased territorial aggression. However, similar to the total body *Tph2* KO mice, the conditional KO animals, exhibited reduced gut motility and decreased anxiety-like behavior. These observations reveal that peripheral serotonergic neurons contribute to control of intestinal motility and anxiety-like behavior and suggest that therapeutics targeting this subset of peripheral neurons could be beneficial.

## Introduction

The central nervous system (CNS) is derived from the neural tube, while the peripheral nervous system (PNS), which includes the dorsal root ganglia, and autonomic and enteric nervous systems (ENS), arises from embryonic neural crest cells (1). While the CNS is critical for higher cognition and behavior, the role of the PNS in these processes is less clear. However, both the PNS and CNS contribute to gastrointestinal physiology, including gastrointestinal motility and secretion (2).

Only a small subset of neurons in the CNS produce 5-hydroxytryptamine (5-HT), the neurotransmitter commonly known as serotonin. In the CNS, serotonergic neurons originate in the brainstem and ramify throughout the brain, regulating a variety of brain functions and behavior through a set of serotonin receptors (3). Although serotonergic neurons in the CNS have been the focus of much research, the production and function of serotonin by neurons in the PNS has received relatively little attention. Recently, single-cell RNA-seq analyses have shown that peripheral serotonergic neurons represent ∼1% of sympathetic and enteric neurons (4, 5). Neuronal serotonin production relies on the enzyme tryptophan hydroxylase 2 (*Tph2*) (6). Mice lacking *Tph2* in all neurons exhibit several behavioral disorders, including decreased anxiety-like behavior and increased territorial aggression as well as reduced gut motility (7, 8). The existence of enteric serotonergic neurons was initially described almost 6 decades ago (9), and they have been implicated to play critical roles in gut motility (7) and potentially depression (10). However, the contribution of serotonergic neurons in the PNS to social behaviors is not well-established. To answer this question, we generated a PNS-conditional knockout mouse (*Tph2*^*fl/fl*^; *Hand2-Cre*), leveraging the *Hand2* promoter, which is active in neural crest-derived cells including enteric neurons and sympathetic neurons in the PNS (11), but not in the brain (12), to selectively investigate the function of serotoninergic neurons in the PNS.

## Results

### Peripheral serotonergic neurons control gut motility

Whole-body *Tph2* KO animals have reduced gut motility and it is thought that the absence of serotonin production from enteric neurons may contribute to this defect (7). To corroborate the importance of peripheral neuronal serotonin in intestinal motility, we analyzed total GI transit time in whole-body and PNS-conditional KO mice. Both *Tph2*^-/-^ and *Tph2*^*fl/fl*^; *Hand2-Cre* mice had diminished total gut motility compared to co-housed littermate controls (Figures 1A). Furthermore, the propulsion of FITC-dextran down the small intestine was significantly slower in *Tph2*^*fl/fl*^; *Hand2-Cre* mice than in their WT littermates (Figures 1B). The results of these two independent assays demonstrate that peripheral serotonergic neurons contribute to intestinal motility. While it is clear that the PNS contributes to gut motility, the impact of CNS serotonin signaling on intestinal motility cannot be determined from these data because of differences in microbiota between the whole-body and PNS-conditional KO animals.

**Figure 1.**
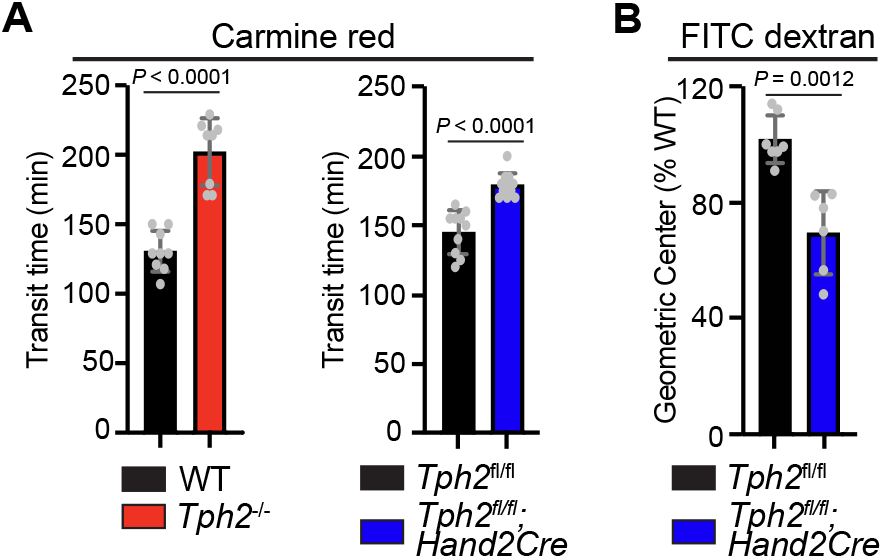
Peripheral serotonergic neurons control gut motility. (A) Total GI transit time was measured using a carmine red assay in WT (n = 9) and *Tph2*^*-/-*^ (n = 8) as well as *Tph2*^*fl/fl*^ (n = 10) and *Tph2*^*fl/fl*^; *Hand2-Cre* (n = 12) mice. (B) FITC-dextran assay of small intestinal motility in *Tph2*^*fl/fl*^ (n = 7) and *Tph2*^*fl/fl*^; *Hand2-Cre* (n = 6) mice. Data shown are means ± SD. Statistical analysis was performed by two-tailed Mann-Whitney test.

### Peripheral serotonergic neurons influence specific aspects of behavior

Our PNS-conditional KO mice enabled us to begin to separate the function of serotonergic neurons in the CNS and PNS in the control of behavior. We assessed behaviors of *Tph2*^*fl/fl*^ and *Tph2*^*fl/fl*^; *Hand2-Cre* mice to investigate whether peripheral serotonergic neurons contribute to the behavioral disorders that have been observed in the whole-body *Tph2* KO mice (8, 13). The absence of neuronal serotonin production in mice and rats lacking the *Tph2* gene is associated with heightened territorial aggression, measured in resident-intruder assays, as well as decreased anxiety-like behavior, measured in elevated plus and elevated zero maze tests. In these experiments, we first replicated reported behavioral defects in *Tph2*^*-/-*^ mice (Figures 2A-2D and movie S1). Co-housed littermates were used for these comparisons in order to control for potential microbiota-related effects.

**Figure 2.**
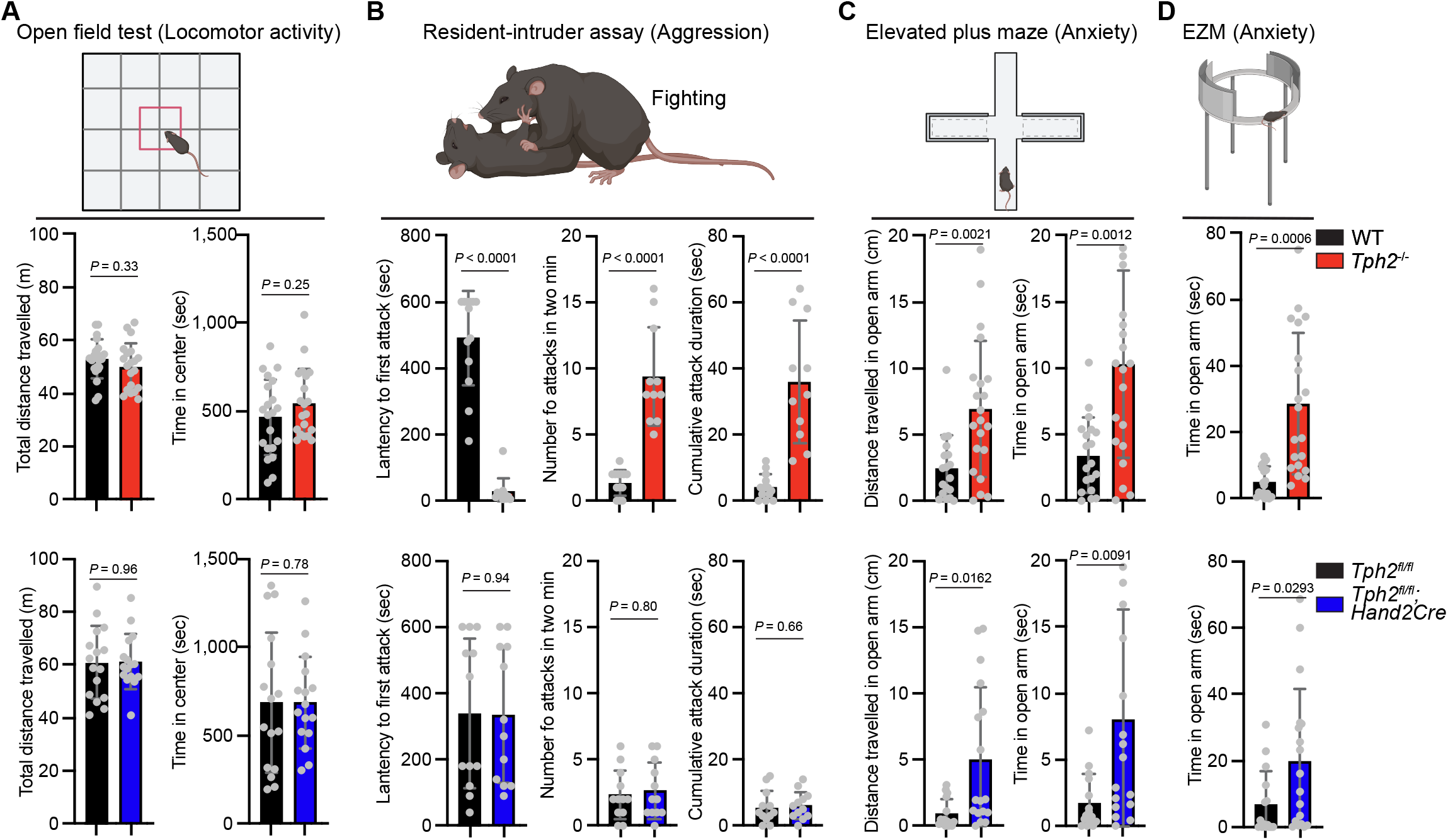
Peripheral serotonergic neurons regulate specific aspects of behavior. (A) Open field test measures of total distance traveled, and time spent in the center area of the open field in mice of the indicated genotypes (WT, n = 21; *Tph2*^*-/-*^, n = 20; *Tph2*^*fl/fl*^, n = 16; and *Tph2*^*fl/fl*^; *Hand2-Cre*, n = 16). (B) Resident-intruder assay measures of the latency to the first attack, number of attack bites and cumulative attack duration by WT (n = 13) and *Tph2*^*-/-*^ (n = 11) as well as *Tph2*^*fl/fl*^ (n = 12) and *Tph2*^*fl/fl*^; *Hand2-Cre* (n = 12) mice. (C) Elevated plus maze assay measures of the total distance traveled, and time spent in the open arms by WT (n = 18) and *Tph2*^*-/-*^ (n = 21) as well as *Tph2*^*fl/fl*^ (n = 15) and *Tph2*^*fl/fl*^; *Hand2-Cre* (n = 17) mice. (D) Elevated zero maze (EZM) measures of time spent in the open arms for WT (n = 13), *Tph2*^*-/-*^ (n = 11), *Tph2*^*fl/fl*^ (n = 12), and *Tph2*^*fl/fl*^; *Hand2-Cre* (n = 12). Data shown are means ± SD. Statistical analysis was performed by two-tailed Mann-Whitney test in A-D.

As observed in whole-body *Tph2* KO animals (8), the deficiency of *Tph2* in the PNS did not impact overall locomotor activity (Figure 2A), suggesting that central and peripheral serotonergic neurons have little influence on general movement. In contrast to the *Tph2*^*-/-*^ animals, the *Tph2*^*fl/fl*^; *Hand2-Cre* mice did not exhibit heightened territorial aggression (Figure 2B and movie S1), consistent with the finding that aggression originates from a subset of *Tph2* neurons in the brain (14). Notably, the PNS-conditional *Tph2* KO mice displayed similar increased exploration in the open arm of the elevated plus and zero maze as observed in the whole-body *Tph2* KO animals (Figures 2C and 2D), suggesting that *Tph2* activity in peripheral neurons plays an important role in inhibiting anxiety-like behaviors. Together, these observations suggest that peripheral serotonergic neurons control important and specific aspects of behavior.

## Discussion

While the roles of CNS serotonergic neurons in cognition and behavior have been well-studied, here, we investigated the roles of peripheral neuron-derived serotonin by creating conditional KO mice that are deficient in *Tph2* in the PNS and not the CNS. Our findings indicate that serotonin produced by peripheral serotonergic neurons impacts diverse processes, including gut motility and anxiety-like behavior, raising the possibility that these phenotypes, along with the intestinal immune deficits observed in the conditional KO (15), are linked.

The overlapping and distinct behavioral patterns in whole-body and PNS-conditional *Tph2* KO animals suggest that, at least in part, serotonergic neurons in the PNS control specific aspects of behavior. Territorial aggression appears to be mediated by CNS serotonergic circuits, since the PNS-conditional KO animals did not exhibit elevated aggression. Surprisingly, the *Tph2*^*-/-*^ and *Tph2*^*fl/fl*^; *Hand2-Cre* animals both showed elevated exploratory behavior that is thought to reflect decreased anxiety. The PNS is heterogeneous and *Tph2* is expressed in a small subset of neural crest derivatives in the enteric nervous system, as well as dorsal root ganglia and autonomic neurons (4). Dissecting the specific roles of subsets of serotonergic neurons in the PNS (e.g., in enteric neurons) as well as potential interactions of peripheral and central serotonergic circuits in regulating anxiety-like behaviors is of interest. This research may shed light on a potential mechanism underpinning the gut-brain axis. Nevertheless, both whole-body and conditional *Tph2* KO animals exhibit reduced anxiety-like behaviors, raising the possibility that peripheral serotonergic neurons contribute to anxiety-like behaviors along with CNS serotonergic circuits. Thus, drugs that block serotonin reuptake (SSRIs) may act, at least in part, by targeting serotonin signaling in the PNS. Engineering SSRIs so that they are unable to penetrate the blood-brain barrier could prevent untoward central nervous system-related side effects of new anxiolytic agents targeting the PNS. Furthermore, defining the cellular networks and mechanisms through which peripheral serotonergic neurons regulate behavior may reveal new targets for pharmacological interventions for anxiety as well as disorders of gut motility.

## Materials and Methods

### Mice

C57BL/6, *Tph2*^*flox/flox*^ mice were purchased from the Jackson Laboratory (Bar Harbor, ME, USA); Swiss Webster (CFW) mice were purchased from the Charles River Laboratories (Wilmington, MA, USA); *Tph2*^*-/-*^ mice were a generous gift from Dr. Gerard Karsenty (Columbia University, NY, USA); *Hand2-Cre* transgenic mice were a generous gift from Ruaidhrí Jackson (Harvard Medical School, MA, USA). *Tph2*^*flox/flox*^ mice were backcrossed to C57BL/6J background for at least six generations. All mice were maintained on a 12-hour light/dark cycle and a standard chow diet at the Harvard Institute of Medicine specific pathogen-free (SPF) animal facility. Animal experiments were performed according to guidelines from the Center for Animal Resources and Comparative Medicine at Harvard Medical School. All protocols and experimental plans were approved by the Brigham and Women’s Hospital Institutional Animal Care and Use Committee (Protocol #2016N000416).

### Gut motility assays

Total gastrointestinal transit time: Carmine red, which cannot be absorbed from the lumen of the gut, was used to study total GI transit time. A solution of carmine red (200 μl; 6%; C1022, Sigma) suspended in 0.5% methylcellulose (M0512, Sigma) was administered by gavage through a 21-gauge round-tip feeding needle. The time at which gavage was carried out was recorded as T0. After gavage, fecal pellets were monitored at 10 min intervals for the presence of carmine red. Total GI transit time was calculated as the interval between T0 and the time of first observance of carmine red in stool.

Small intestine transit time: A solution containing fluorescein isothiocyanate-dextran (FITC-Dextran, 100 μl; 10 mg/ml in 2% methylcellulose; FD70, Sigma) was administered to mice by gavage through a 21-gauge round-tip feeding needle. Animals were sacrificed 15 min after gavage; the small intestine, cecum, and colon were collected in PBS. The small intestine was divided into 10 segments of equal length, and the colon was divided in half. Each piece of tissue was transferred into a 2 ml tube containing 1 ml of PBS, homogenized, and centrifuged (2000 × g), yielding a clear supernatant. FITC fluorescence was measured in the supernatant (Epoch2, Biotek). Small intestinal transit was estimated by the position of the geometric center of the FITC-dextran in the small bowel. For each segment of the small intestine (1-10), the geometric center (a) was calculated as follows: a = (fluorescence in each segment × number of the segment)/(total fluorescence recovered in the small intestine). The total geometric center is Σ (a of each segment) and was normalized to WT littermates.

### Mouse behavior assays

The following assays were carried at the Harvard Medical School Mouse Behavior Core according to standard methods by an observer blinded to the genotype of the animal:

#### Open Field

a large arena (50 × 50 cm) under low illumination (30 lux) was used as an open field to measure locomotor activity. Each mouse was placed into the arena and its activity was measured during 10 min. The total distance traveled, and time spent in the center were calculated.

#### Resident-intruder assay

This assay was performed largely as described(14). Transgenic male mice, ‘‘residents’’ were group-housed with male siblings until sexual maturity. Resident males were assayed for aggression toward a wild-type Swiss Webster (CFW) (Charles River) intruder mouse. Each intruder mouse was used only once to avoid submissive/dominance effects after the initial interaction. Behavioral interactions during each confrontation were recorded and subsequently scored by an observer blinded to the mouse genotype. Latency to the first attack, total amount of attacks and cumulative duration of attacks were analyzed.

#### Elevated plus maze (EPM) and zero maze (EZM)

The EPM and EZM test is based on the aversion of rodents to open, bright illuminated spaces. The EPM consisted of two open arms (30 × 5 cm) and two closed arms (30 × 5 cm) that were enclosed by a sidewall on all edges (height 15cm). The EZM consisted of a gray plastic annular runway (width 6 cm, outer diameter 46 cm, 50 cm above ground level). Two opposing sectors were protected by inner and outer walls of gray polyvinyl (height 15 cm). Mice were placed in one of the closed arms of the maze. Distance travelled and time spent in open arms were quantified during 10 min tests. Arm entry was only scored when an animal (the mouse mass center) was at least 3 cm in an arm

## Statistical methods

Statistical analyses were carried out using the two-tailed Student’s *t*-test, two-tailed Mann-Whitney test or Kaplan-Meier Log-rank test on GraphPad Prism5 (version 9.4.1).

## Supporting information

Supplemental vedio

## Acknowledgments

We thank members of the Waldor lab and Drs. Brandon Sit and Susan M. Dymecki for helpful discussions on all aspects of this project, Dr. Gerard Karsenty at Columbia University for *Tph2*^*-/-*^ mice, and Dr. Olga Alekseenko and Dr. Barbara Caldarone at the Harvard Mouse Behavior Core for assistance with mouse behavioral testing.

Research in the M.K.W. laboratory is supported by HHMI and NIH grant R01 AI-042347.

## License information

This article is subject to HHMI’s Open Access to Publications policy. HHMI lab heads have previously granted a nonexclusive CC BY 4.0 license to the public and a sublicensable license to HHMI in their research articles. Pursuant to those licenses, the author-accepted manuscript of this article can be made freely available under a CC BY 4.0 license immediately upon publication.

## Figures

**Supporting movie S1**. Analyzing aggressive behavior via the resident-intruder assay in *Tph2*^*-/-*^ (left panel) and *Tph2*^*fl/fl*^; *Hand2-Cre* (right panel) mice.

