## Supplemental figures for "Peripheral serotonergic neurons regulate gut motility and anxiety-like behavior": Fig S1.pdf

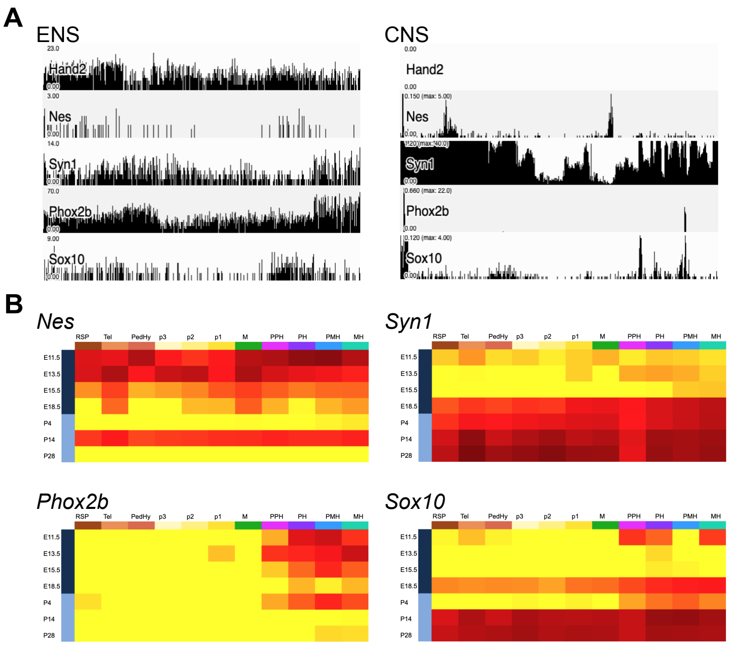


(A) In the mouse brain neuron database at Mousebrain.org.level6/L6_Cns_neurons.loom, the gene expression of *Nes*, *Syn1, Phox2b, Sox10* (commonly used cre-driver for enteric neurons) is evident both in ENS and CNS. However, the presence of *Hand2* gene expression in CNS is not detected.

(B) In the Allen Developing Mouse Brain Atlas database, gene expression was observed for *Nes*, *Syn1*, *Phox2b,* and *Sox10* (a commonly used cre-driver for enteric neurons) in the CNS. However, *Hand2* gene expression in the CNS was not detected.

**Figure S1**
